# Predicting brain functions from structural connectome using graph neural network

**DOI:** 10.1101/2022.03.31.484925

**Authors:** Edward S. Hui, Yuxiang Sun, Ho Ko, Chetwyn C.H. Chan, Peng Cao

**Affiliations:** Department of Rehabilitation Sciences, The Hong Kong Polytechnic University, HKSAR, China; Department of Mechnical Engineering, The Hong Kong Polytechnic University, HKSAR, China; Division of Neurology, Department of Medicine and Therapeutics, The Chinese University of Hong Kong, HKSAR, China; Department of Psychology, The Education University of Hong Kong, HKSAR, China; Department of Diagnostic Radiology, The University of Hong Kong, HKSAR, China

**Keywords:** Structural connectome, prediction of brain functions, graph neural network, attention mechanism

## Abstract

The relationship between brain structure and function remains elusive, amidst the tremendous advances in brain mapping techniques. In this work, we attempt to partially disentangle this relationship by connecting task–evoked functional MRI (fMRI) responses with the underlying structural connectome using graph neural network (GNN). MRI data (n = 1,063) were collected from the Human Connectome Project. We demonstrate that our GNN–based model predicts task–evoked fMRI responses with high fidelity. Using a graph attention mechanism, it is possible to infer the subsets of neighboring cortical regions whose structural connections are important for the prediction of the functional responses of individual cortical regions. Notably, for each cortical region, such subset of neighboring cortical regions is predominantly localized to the ipsilateral hemisphere and much smaller than that with direct structural connections. We found that the higher cognitive functions subserved by the cingulo–opercular, dorsal attention, frontoparietal and default mode clusters may depend on neighboring cortical regions across a wide range of functional brain clusters in the ipsilateral hemisphere, whilst the sensory functions subserved by the visual1 and auditory clusters on neighboring cortical regions across much fewer functional brain clusters.

## 1 INTRODUCTION

The advent of imaging technologies has permitted detailed and comprehensive mapping of the structure and function of human brain [1, 2, 3]. However, the critical knowledge on how brain function emerges from structure remains elusive.

That cognition is supported by large-scale interactions between different brain regions [4] suggests that the connectomics approach, the study of anatomical and functional connections across the whole brain [4, 5], may shed new light on the elusive brain structure–and–function relationship. Several models have been proposed to bridge the link between brain function and connectome, including statistical [6, 7, 8, 9, 10, 11, 12], network communication [13, 14], and machine learning model [15, 16, 17, 18, 19, 20].

Motivated by the notion that dynamic functional brain connections are largely facilitated and constrained by the underlying static brain anatomical connections [21], another potential avenue toward elucidating the elusive brain structure–and–function relationship may pertain to interrogating the anatomical connections that underpin brain functions. Thiebaut de Schotten et al. [11] and Nozais et al. [12] have both attempted to map the task–evoked functional MRI (fMRI) responses of brain anatomical connections. Notwithstanding these instrumental efforts in improving our understanding on the brain structure–and–function relationship, two questions regarding the brain structure–and–function relationship remain unexplored. On which subset of neighboring brain regions do the functions of a given brain region depend? How would this “dependence” differ between brain regions from different functional brain clusters?

In this study, we attempt to partially address these questions by developing a graph neural network [22] model that is able to predict the task–evoked fMRI responses of individual cortical regions from structural connectome. We hypothesize that our model could partly disentangle the relationship between brain functions and structure by virtue of inferring the structural connections most important for predicting the task–evoked fMRI responses individual cortical regions.

## 2 METHODS

### 2.1 Datasets

Data from normal twin and non-twin siblings (*n* = 1, 063; ages: 22 − 35 years) were obtained from the Human Connectome Project (HCP) [1]. The minimally pre–processed MRI data and task–based fMRI analysis results were used (see ref. [3] for details on the pre– and post–processing pipelines).

### 2.2 Construction of connectome and brain activation

The HCP pipelines [3], FMRIB’s software library 6.0.4 (FSL) and Connectome Workbench 1.4.2 were used. The fiber orientation density function was estimated from diffusion MRI data using FSL’s BED- POSTX (`bedpostx_gpu -n 3 -b 3000 -model 3 -g` --`rician`). A dense connectome containing the number of fiber tracks between a pair of grayordinates was subsequently obtained using probabilistic tractography using FSL’s PROBTRACKX (`probtrackx2_gpu` --`loopcheck` –`forcedir` --`fibthresh=1 -c 0.2` --`sampvox=2` --`randfib=1 -P 3000 -S 2000` --`steplength=0.5 -s -m` --`meshspace=caret -x` --`seedref` --`xfm` --`invxfm` --`stop` --`wtstop` –`waypoints` --`omatrix3` --`target3`). The dense connectome was then parcellated using a brain parcellation with 360 cortical regions by Ji et al. [23] using Connectome Workbench (`wb_command -cifti-parcellate -only-numeric`).

Whole-brain activation maps from 47 contrasts (see Supplementary Table S1 for full list) from emotion, gambling, language, motor, relational, social and working memory tasks were obtained from the task–based fMRI analysis results (see ref. [24] for task details). These activation maps were subsequently parcellated using the same brain parcellation [23] as that of the parcellated structural connectome.

### 2.3 Graph neural network model

Motivated by the fact that static brain anatomical connections may provide an important window for disentangling the complex relationship between brain structure and functions, we propose a graph neural network (GNN) [22] model that can not only integrate the wealth of information embedded in structural connectome for the prediction of brain functions in a nonlinear and principled way, but also permits inference on the structural connections most important for predicting the task–evoked fMRI responses of individual brain regions.

Graph neural network was proposed as an extension to convolutional neural network (CNN), the most popular deep learning model [25], to exploit the topological information inherent in input data that can be represented as a network, as characterized by network nodes and edges (brain is a prime example), to improve model performance [22]. The premise of GNN is that the features of a network node or brain region *v* (denoted as target cortical region from hereon) **z**_*v*_ ∈ ℝ^*F*′^, where *F*′ is the number of features at output, can be learnt from the features of *v*’s neighboring nodes or brain regions *u* ∈ 𝒩(*v*) (denoted as source cortical regions from hereon) **x**_*u*_ ∈ ℝ^*F*^, where *F* is the number of input features and 𝒩(*v*) the neighborhood of node *v*. For our proposed GNN–based model, the task–evoked fMRI responses of a target cortical region were learnt from the structural connectome of its source cortical regions. In other words, the parcellated task–evoked responses of the entire cortex **Z** = [**z**_1_ … **z**_*N*_]^T^ ∈ ℝ^*N* ×*F*′^, where *N* is the number of cortical regions, were learnt from the underlying structural connectome **A**_0_ ∈ ℝ^*N* ×*N*^ (see below for the exact relationship between **A**_0_ and **X** = [**x**_1_ … **x**_*N*_]^T^ ∈ R^*N* ×*F*^). For the brain parcellation by Ji et al. [23], *N* = 360. The task–evoked fMRI responses were obtained from *F*′ = 47 contrasts from 7 different cognitive tasks.

The detailed network architecture of our GNN-based model is shown in Figure 1. The core innovation of our model is that it is interpretable, thereby permitting the inference on the subset of source cortical regions whose structural connectome are important for the prediction of the task–evoked fMRI responses of a target cortical region via the attention block (ATT; the left side of Figure 1), which is based on multi-head graph attention network (GAT) [26]. The premise of GAT is that, instead of applying equal weight to the features of source cortical regions in the process of model training, these weights, also known as attention weights **Φ** ∈ ℝ^*N* ×*N*^, depend on a trainable combination of the features of the target cortical region and the features of its source cortical regions (i.e., 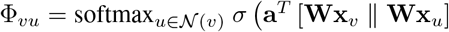, where *σ* (·) is activation function, **a** ∈ ℝ^2*F*′^ and **W** ∈ R^*F*′ ×*F*^ are trainable parameters, and ∥ concatenation operation) [26]. Because of these region-dependent **Φ**, the subset of source cortical regions most important for the prediction of the task–evoked fMRI responses of a given target brain region can be subsequently inferred. To increase model capacity and performance, we have adopted a multibranch approach, whereby there were *N*_*i*_ GNN–based blocks (dotted rectangles in Figure 1) in different parts of the network, with each receiving **Â**_0_ subjected to different trainable threshold 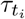 (i.e., 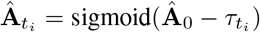, where 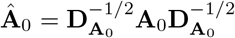 and 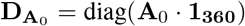). As such,the input features **X** to our GNN-based model at the ATT at a given 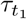 were subsequently obtained by row–normalizing 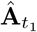 (i.e., 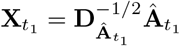).

**Figure 1:**
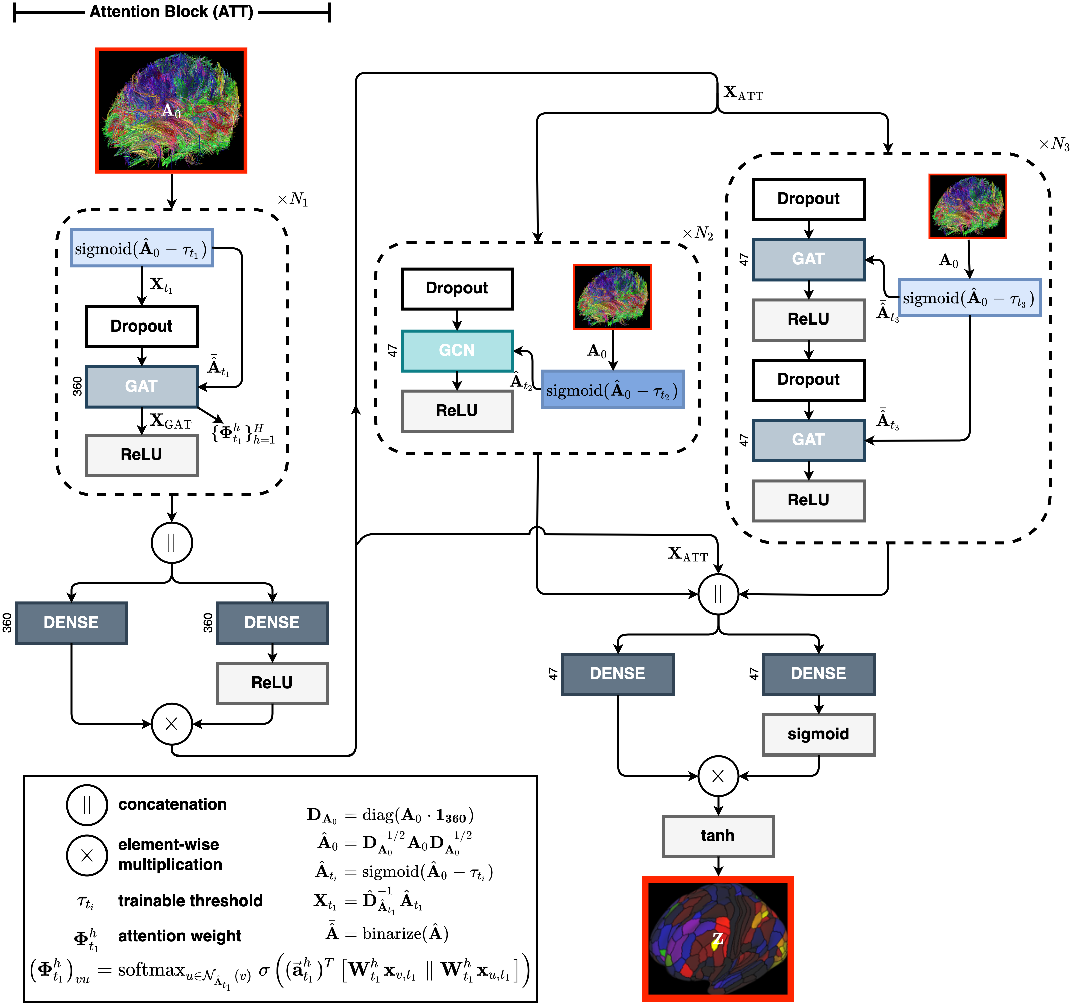
The architecture of our GNN–based model for the prediction of task–evoked functional MRI responses of a target cortical region from the structural connectome **A**_0_. The left half of the model consists of an attention block (ATT), which is based on GAT, that permits inference on the structural connections of source cortical regions most important for the prediction of the task–evoked fMRI responses of target cortical region **Z**. The second half of the model consists of GCN and GAT blocks. Note that the input features 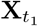 in ATT were the structural connections between target and source cortical regions. *σ* (·) represents activation function, **a** ∈ ℝ^2*F*′^and **W** ∈ ℝ^*F*′ ×*F*^ trainable parameters of GAT,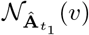 the neighbor- hood of node *v* in the thresholded structural connectome 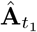. Note also that the structural connectome weights were obtained from 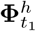 (i.e., 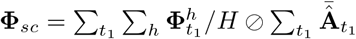), where ⊘ is elementwise division.

### 2.4 Model training

Only the data from subjects whom have completed all task–evoked fMRI experiments were used (*n* = 966). Of these data, 80% were used for training, 15% for validation and 5% for testing. Each of the 47 parcellated task–evoked fMRI responses was normalized by its maximum. Our GNN–based model was trained using the following parameters: ADAM optimizer with learning rates = 0.001, number of epochs = 600 (with early stopping with patience = 30), mini–batch size = 5, number of trainable thresholds for the ATT (*N*_1_), GCN (*N*_2_) and GAT (*N*_3_) blocks = 8, 8, 8, *H* = 8, dropout rate = 0.5, and mean square error as loss function. All models were implemented using Tensorflow v2.4.1. and the spektral [27] library with a workstation with 40-core Intel Xeon 5218R CPU, NVIDIA GV100 GPU with 32GB of memory and 512GB RAM.

### 2.5 Structural connectome weights Φ_*sc*_

A “summary” attention weight matrix **Φ**_*sc*_ ∈ ℝ^*N* ×*N*^, dubbed structural connectome weights from hereon, that permits inference on the subset of source cortical regions (in this case, structural connectome) most important for the prediction of the task–evoked fMRI responses of a target cortical region was estimated from the attention weights from the GAT in the ATT using the following procedures: (1) Since there were *H* heads for each GAT, the attention weights of the *h*^*th*^ GAT head for the 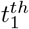 threshold on 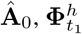, were averaged 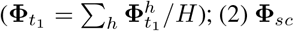 was then obtained by taking an average of all 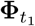 (i.e., 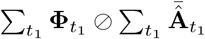, where 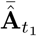 represents unweighted 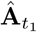, and ⊘ elementwise division). (**Φ**_*sc*_)_*vu*_, where (·)_*vu*_ represents the *vu*^*th*^ element of a matrix, thus approximates the overall weight on the features of node *u* from which the ATT predicts the feature of node *v* at output, **x**_ATT,*v*_. The structural connectome weights at the level of individual unilateral brain clusters 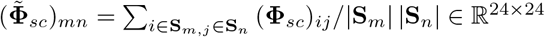, where (·)_*ij*_ represents the *ij*^*th*^ element of a matrix, **S**_*m*_ the set of brain regions in the *m*^*th*^ cluster in one hemisphere, and |**S**_*m*_| the size of **S**_*m*_) were also obtained by using the 12 functional brain clusters of the parcellation by Ji et al. [23].

Subject-specific structural connectome weights 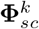 were obtained from the trained model using the *n* = 194 data from the validation and test sets. One-sample t-test was performed to determine whether the structural connectome weight between the *v*^*th*^ target cortical region and the *u*^*th*^ source cortical region (i.e., 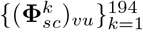) was significantly different from zero. All *p–values* were FDR–corrected for multiple comparisons and analyzed for significance at *α* = 0.05. Subject-averaged structural connectome weights were subsequently obtained (denoted as 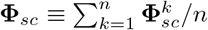 from hereon), with individual weight being zeroed if its corresponding *p-value* ≥ 0.05 after FDR–correction.

The subset of source cortical regions *u* most important for the prediction of the task–evoked fMRI responses of the target cortical region *v* was subsequently identified from the entries of **Φ**_*sc*_ that were larger than zero (i.e., {*u* : (**Φ**_*sc*_)_*vu*_ > 0 ∀ *u* ∈ 𝒩(*v*)}).

### 2.6 Evaluation of model performance

The performance of our GNN-based model was evaluated using Pearson correlation between the task–evoked fMRI responses of each functional brain cluster 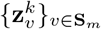 from ground truth versus prediction for each of the *n* = 49 subjects in the test set. Significance was set at *p* < 0.05.

## 3 RESULTS

### 3.1 Model performance

The ground truth and model prediction of the parcellated task–evoked fMRI responses of two representative subjects are shown in Figure 2a. Pearson correlation between the task–evoked fMRI responses of individual functional brain clusters from the ground truth versus model prediction for each subject in the test set were also performed (see Figure 2b). The model prediction was largely consistent with the ground truth, with the visual1 and visual2 clusters having the strongest correlation and the orbito–affective cluster the lowest.

**Figure 2:**
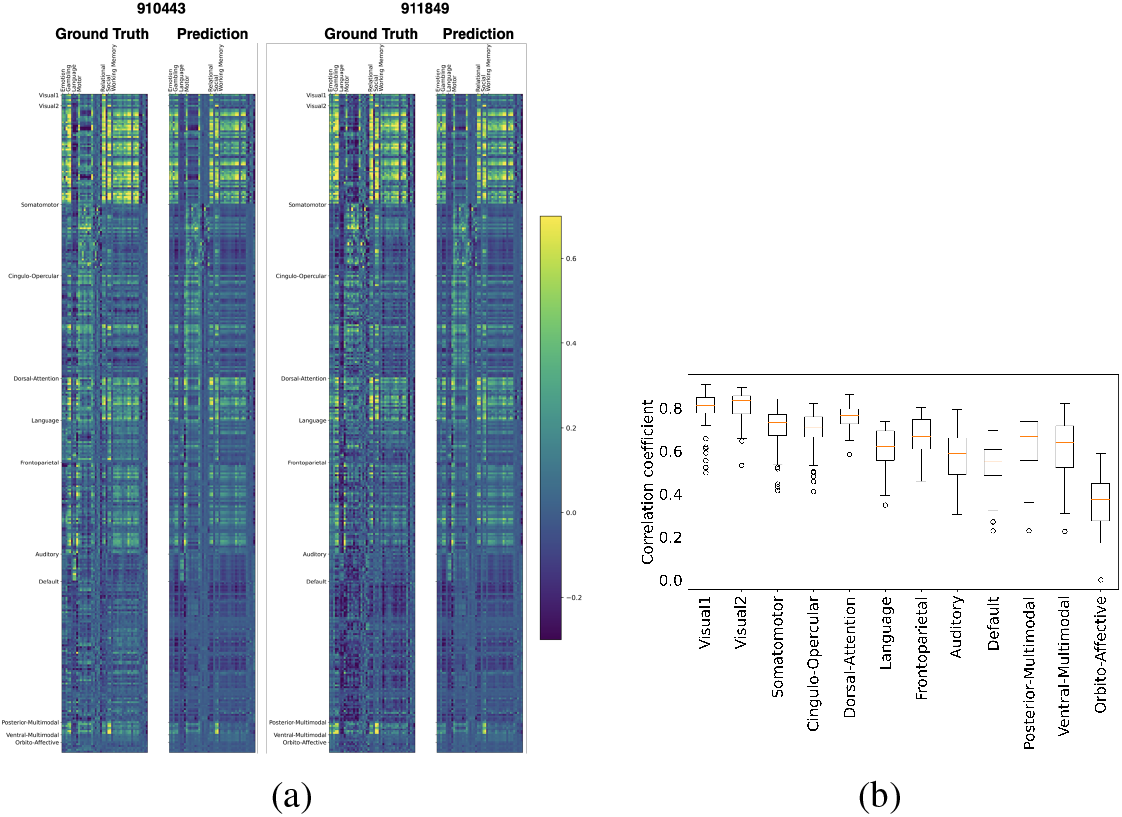
(a) The true and predicted parcellated task–evoked fMRI responses from 47 contrasts of 2 representative subjects (ID: 910443 and 911849). (b) The Pearson correlation between the task–evoked fMRI responses of individual functional brain clusters from ground truth and model prediction for each of the *n* = 49 subjects from the test set.

### 3.2 Structural connectome weights

The subject–averaged **Φ**_*sc*_ is shown in Figure 3a (only the entries with FDR-corrected *p* < 0.05 are shown). Evident from Figure 3a, **Φ**_*sc*_ was much sparser than the input structural connectome **Â**_0_, suggesting that not all source cortical regions are required for predicting the task–evoked fMRI responses of target cortical regions. The characteristics of **Φ**_*sc*_ and **Â**_0_ at the level of individual functional brain clusters are respectively summarized in Table 1. Compared to **Â**_0_, the cluster–averaged degree of source regions of **Φ**_*sc*_ were substantially smaller (third column of Table 1). The number of clusters to which source regions belong was smaller (last column of Table 1), and the percent of source regions residing in the same functional brain cluster as that of the target regions were higher (underlined in Table 1).

**Table 1:**
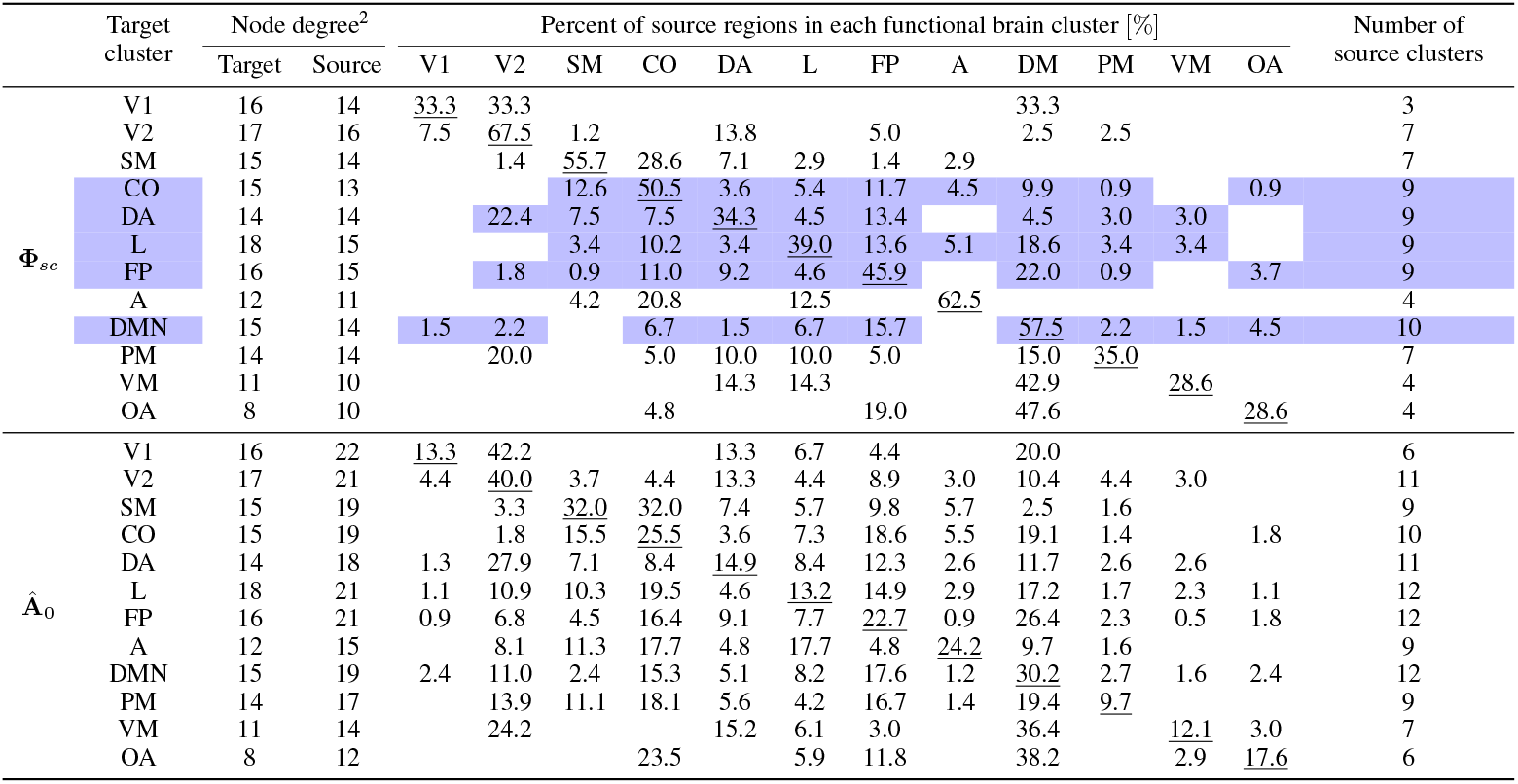
Characteristics of target cortical regions at the level of functional brain cluster for **Φ**_*sc*_ and **Â**_0_ from *n* = 194 data from the validation and test sets. The subset of source cortical regions *u* most important for the prediction of the task–evoked fMRI responses of the target cortical region *v* was identified from {*u* : (**Φ**_*sc*_)_*vu*_ > 0 ∀ *u* ∈ 𝒩(*v*)}. V1: visual1; V2: visual2; SM: somatomotor; CO: cingulo–opercular; DA: dorsal attention; L: language; FP: frontoparietal; A: auditory; DM: default mode; PM: posterior–multimodal; VM: ventral–multimodal; OA: orbito–affective.

**Figure 3:**
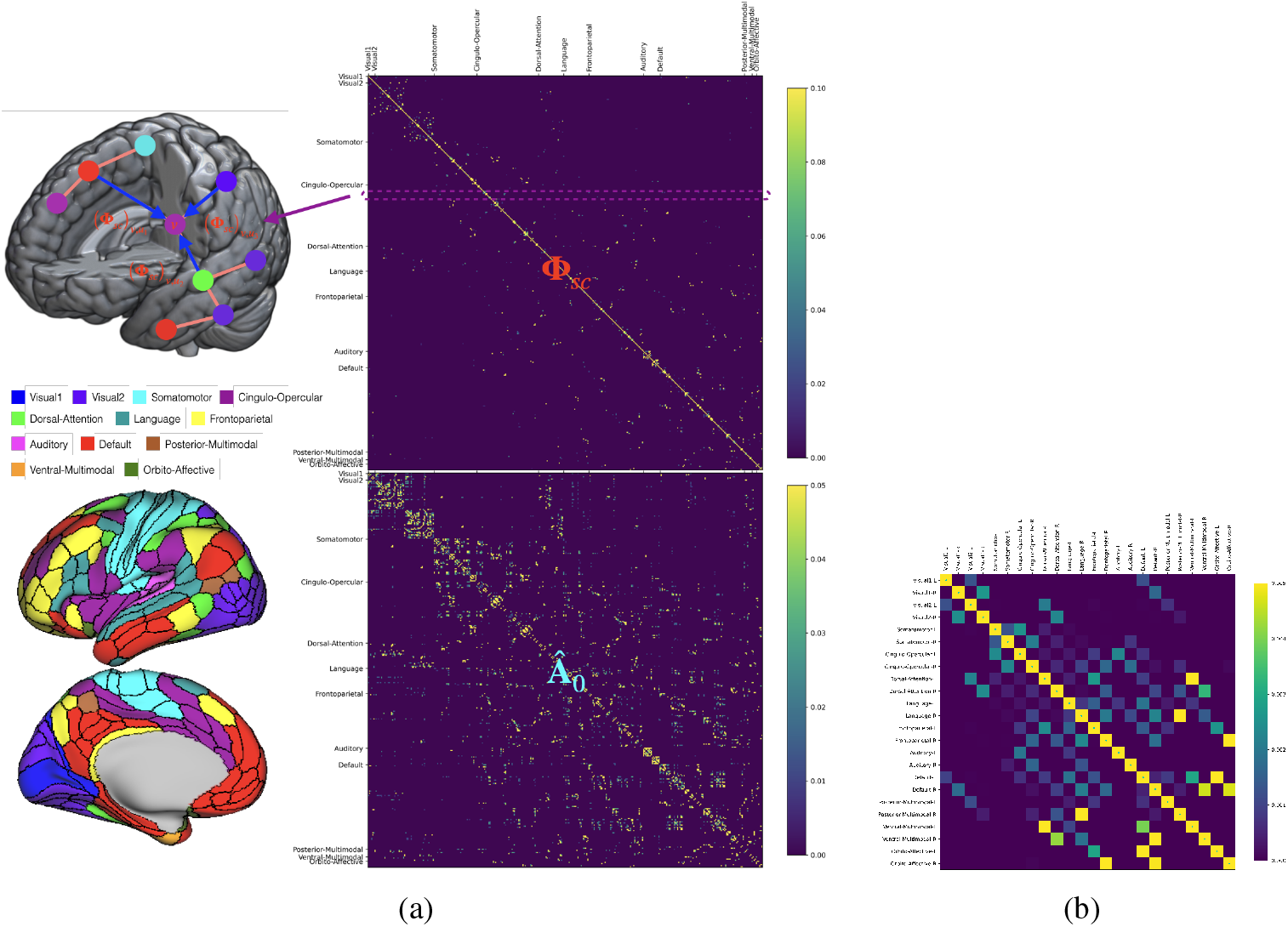
The inference on the subset of source cortical regions whose structural connectome are important for the prediction of the task–evoked fMRI responses of a target cortical region was made possible by the structural connectome weights **Φ**_*sc*_ (top right in (a); target regions were arranged in different rows, and source regions in different columns; only the entries with FDR-corrected *p* < 0.05 were displayed). (**Φ**_*sc*_)_*vu*_ approximates the overall weight on the features of node *u* from which ATT predicts the feature of node *v* at output, **x**_ATT,*v*_. Comparing **Φ**_*sc*_ with the input structural connectome **Â**_0_ (bottom right in (a)), it is evident that the structural connectome of only a small subset of source cortical regions were required for predicting the task–evoked fMRI responses. (b) The structural connectome weights at the level of functional brain cluster in each hemisphere 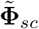.

The checkerboard pattern apparent in 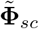 (see Figure 3b) suggests that most of the target and source cortical regions were ipsilateral to each other. Target regions were thus dichotomized according to whether the prediction of their task–evoked fMRI responses depended on the structural connectomes of source regions in the ipsilateral hemisphere or both hemispheres. The cluster–averaged characteristics of **Φ**_*sc*_ and **Â**_0_ for these two groups of target cortical regions are summarized in Tables 2. Compared to **Â**_0_, the task–evoked fMRI responses of the majority of target regions of **Φ**_*sc*_ depended on the structural connections of ipsilateral source regions (c.f. the count in the second column of Table 2). For target regions whose task–evoked fMRI responses depended on the structural connections of source regions from bilateral hemispheres (somatomotor: 23%, cingulo-opercula: 4%, language: 9%, frontoparietal: 4%, and default mode: 13%), majority of the source regions belonged to the same functional brain cluster of that of the target regions.

**Table 2:**
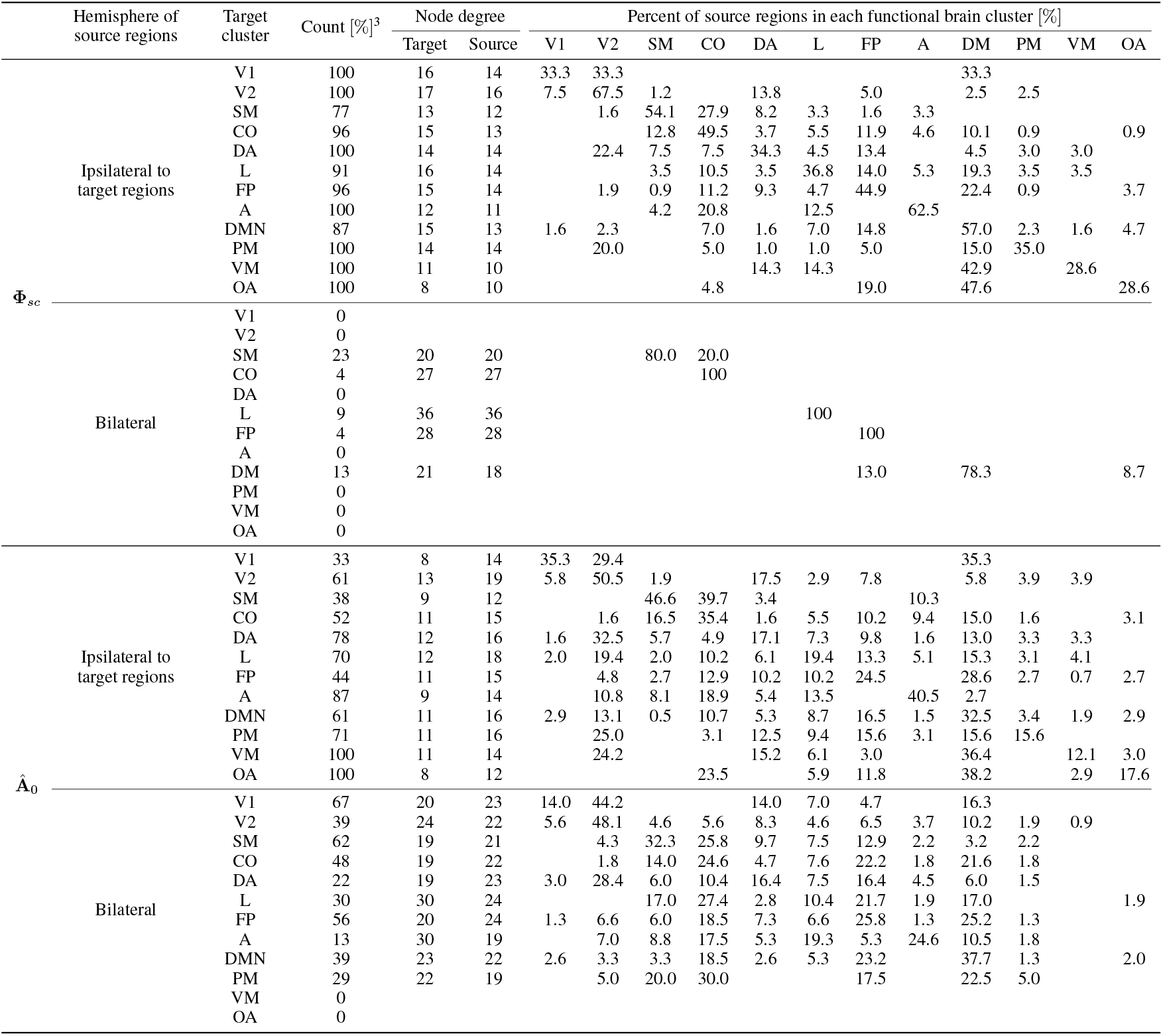
Cluster-averaged characteristics of a subset of target cortical regions of **Φ**_*sc*_ and **Â**_0_ from *n* = 194 data from the validation and test sets.

## 4 DISCUSSION

Considering that the intercommunication between different parts of the brain elicited by cognition and behavior should depend on the underlying structural connections, numerous studies have exploited this fact to model the elusive brain– structure–function relationship. Zimmermann et al. modelled the relationship between cognitive trait and structural connections using partial least square [28]. Ekstrand et al. predicted the voxel–wise task–evoked responses from structural connectome using a linear regression model [29]. Liu et al. modelled the relationship between structural connectome and cognitive traits using a nonlinear latent factor model that utilized regression and graph auto–encoder [30].

Our GNN–based model on the other hand aims to further disentangle the brain structure–and–function relationship by inferring the subset of source cortical regions most important for the prediction of the task–evoked fMRI responses of individual target cortical regions.

### 4.1 Toward the neuroarchitectural underpinning of brain functions

The fact that **Φ**_*sc*_ was much sparser than **Â**_0_ (c.f., Figure 3a) and the cluster–averaged degree of the source cortical regions of **Φ**_*sc*_ was smaller than those of **Â**_0_ (c.f., the third column of Table 1) suggest that the task–evoked fMRI responses of a target region might depend on the structural connections of a subset of its neighboring cortical regions that were subsequently connected to fewer number of cortical regions. These source cortical regions were distributed across a wide range of functional brain clusters for target regions in the cingulo–opercular, dorsal attention, language, frontoparietal and default mode clusters (c.f., highlighted in blue in Table 1), and across much fewer clusters for target regions in the visual1, auditory, ventral–multimodal and orbito–affective clusters (also see Figure 4 for an overview on this cluster–averaged target–source–dependence). Furthermore, majority of these source regions were ipsilateral to target regions, whilst the rest were distributed across the somatomotor, cingulo–opercular, language, frontoparietal and default mode clusters in both hemispheres (c.f., Table 2).

**Figure 4:**
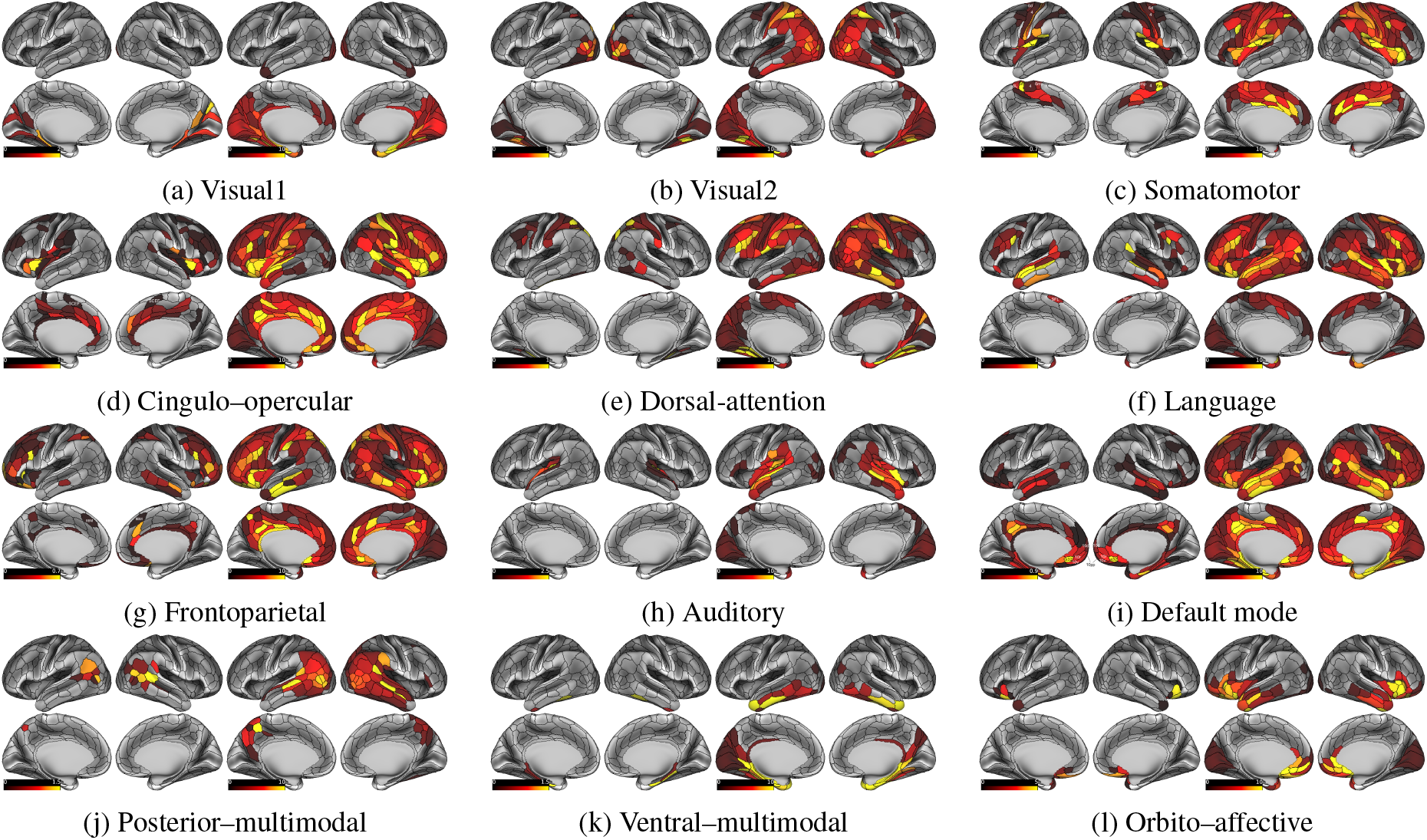
Rendering of the cluster–averaged 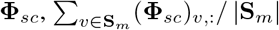, where (·)_*v*,:_ is the *v*^*th*^ row of a matrix, are illustrated in the first and second columns of each subfigure. The cluster-averaged **Â**_0_ of source cortical regions, 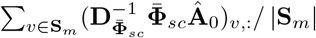, where 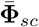 represents unweighted **Φ**_*sc*_, which illustrates the cortical regions to which source cortical regions are connected, are also illustrated in the third and forth columns of each subfigure.

Taken together, these findings suggest that the higher cognitive functions subserved by the cognitive systems, such as the cingulo–opercular, dorsal attention, frontoparietal and default mode clusters, may require the “support” from source cortical regions across a wide range of functional brain clusters in the ipsilateral hemisphere, whilst the sensory functions subserved by the sensory systems, such as the visual1 and auditory clusters, from source cortical regions across much fewer functional brain clusters. This difference in the “scope” of support from source cortical regions may underpin the difference in the effect of sensory cues versus cognitive cues on visual attention, a mechanism for filtering behaviorally relevant input. It was previously reported that sensory cues permit more rapid facilitation of the detection and discrimination of input than cognitive cues [31]. This may suggest that the cortical regions in the sensory systems process and relay information faster than those in the cognitive systems. Supported by our results, one plausible mechanism underlying this difference in processing speed may pertain to the functions of sensory systems being supported by fewer functional brain clusters (thus possibly requiring less time to process information) than those of the cognitive systems.

Our results also show that the task–evoked fMRI responses of the majority of target cortical regions depended on ipsilateral source cortical regions, whilst a small number of target regions in the somatomotor, cingulo–opercular, language, frontoparietal and default mode clusters depended on source regions in the contralateral hemisphere or both hemispheres (listed in Table 3, and labeled in Figures 4d, f, g, i). Such “lateralization” coincides with the functional asymmetry between the two hemispheres (a.k.a. hemispheric lateralization) that was posited to be due to diminished reliance on long-range interhemispheric communications [32]. Our findings also partly corroborates with those of Zimmermann et al., wherein significant brain–structure–cognition relationships were mostly found in the structural connections between ipsilateral brain regions (see Figure 3 in ref. [28]).

**Table 3:**
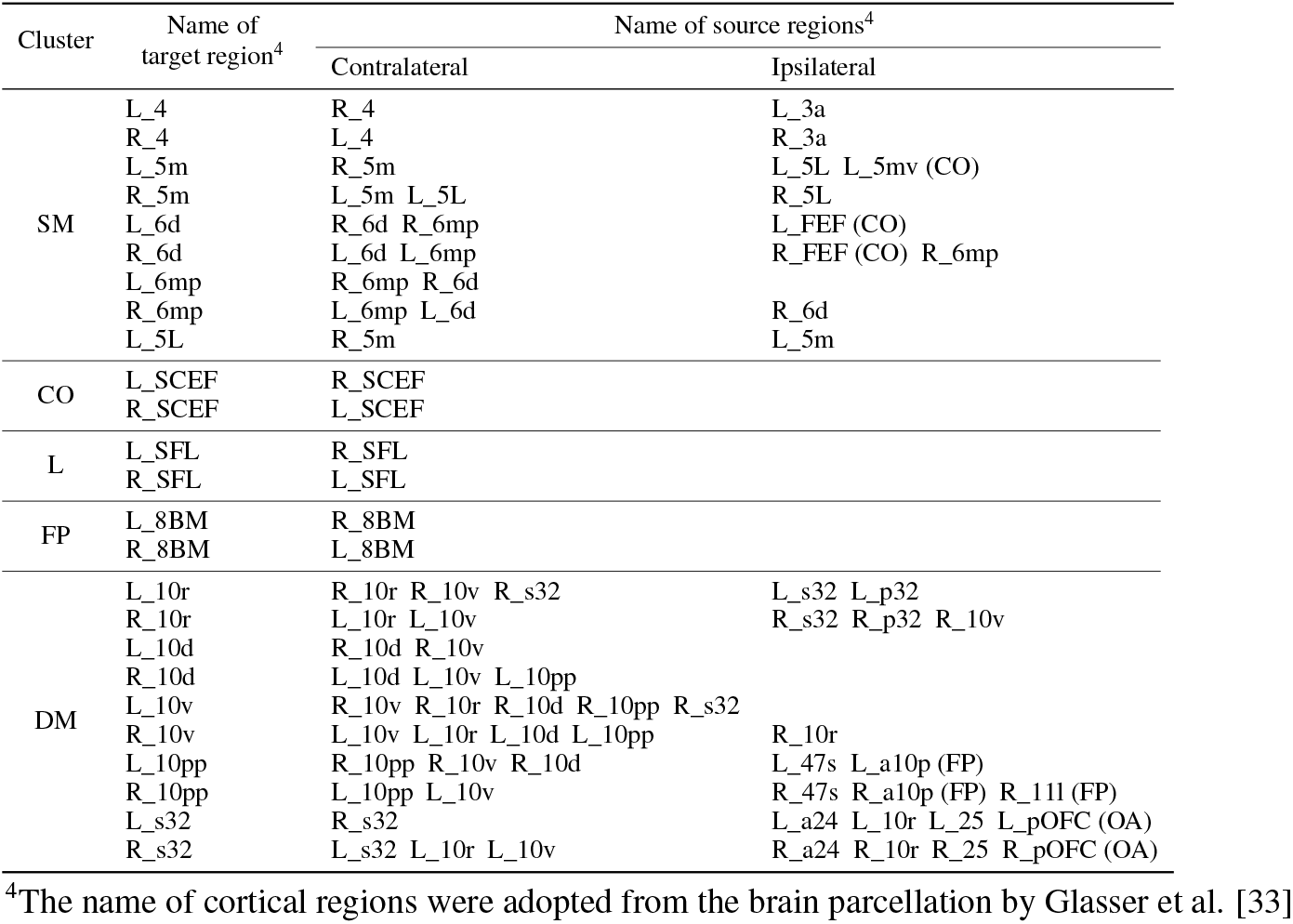
Target cortical regions whose functions were supported by source cortical regions in the contralateral hemisphere. L_: left hemispheric; R_: right hemispheric; FEF: frontal eye field; SCEF: supplementary and cingulate eye fields; SFL: superior frontal language; pOFC: posterior orbitofrontal complex.

### 4.2 Limitations and future studies

A major limitation of our GNN-based model is that the subject-specific variations in task–evoked fMRI responses were not fully captured by our model, rendering similar model predictions for different subjects (see Figure 2a). In future study, we will investigate if this problem can be alleviated with larger training sets. Furthermore, although the structural connectome weights permitted inference on the subset of source regions most important for model prediction, it was not possible to rank these source regions to identify the dominant component for prediction.

There are a number of modifications to our GNN-based models that remain to be investigated. First, apart from structural connectome, other imaging biomarkers, such as the cortical thickness, myelin content, diffusion metrics, could also be incorporated in the input feature matrix. Second, all 7 distinct types of behavioral tasks were incorporated in the output feature matrix. It remains to be investigated how would structural connectome weights change for each type of tasks. Third, we were not able to predict the task–evoked fMRI responses of individual grayordinate due to limited GPU memory. It is worthwhile to investigate the performance of our GNN-based model for such a large brain network (with approximately 60, 000 grayordinates as individual network node).

## 5 CONCLUSION

We have demonstrated that parcellated task–evoked fMRI responses can be predicted from the structural connectome of the entire cortex using graph neural network. We have also attempted to partially disentangle the elusive relationship between brain structure and function by inferring the subset of neighboring cortical regions whose structural connections are important for predicting the task–evoked fMRI responses of individual cortical regions using graph attention network.

---

**Supplementary Table S1:** The name of all 47 contrasts from 7 behavioral tasks. Please refer to ref. [24] for more details on the HCP tasks.

| Task | Contrasts |
| --- | --- |
| Emotion | Faces shapes faces-shapes |
| Gambling | Punish reward punish-reward |
| Language | Math story math-story |
| Motor | Cue LF LH RF RH T AVG Cue-AVG LF-AVG LH-AVG RF-AVG RH-AVG T-AVG |
| Relational | Match REL Match-REL |
| Social | Random TOM Random-TOM |
| Working memory | 2BK_body 2BK_face 2BK_place 2BK_tool 0BK_body 0BK_face 0BK_place 0BK_tool<br>2BK 0BK 2BK-0BK body face place tool body-AVG face-AVG place-AVG tool-AVG |

## Footnotes

2 Average of the degree of cortical regions in the same cluster (i.e.,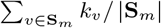, where 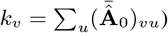)

3 Percent of the cortical regions in target cluster that belong to a subset of target nodes

## Notes

### Competing Interest Statement

The authors have declared no competing interest.

